# Assessing the environmental impacts of conventional and organic scenarios of rainbow trout farming in France

**DOI:** 10.1101/2023.09.15.557864

**Authors:** Simon Pouil, Mathieu Besson, Florence Phocas, Joël Aubin

## Abstract

In France, rainbow trout (*Oncorhynchus mykiss*) farming traditionally used flow-through systems, which raised concerns about environmental impacts, including limited freshwater availability, and the use of ingredients from intensive agriculture and fishing. To address the growing demand for sustainable food products, there is an increasing interest in organic aquaculture. In this study, we employed an attributional life cycle assessment (LCA) to analyse the environmental impacts of rainbow trout production. We simulated conventional and organic production practices in a hypothetical fish farm to evaluate the differences in environmental impacts at the farm level. The potential impacts were calculated using a product-based functional unit (one tonne of trout) under the two production scenarios and were also expressed using a surface-based functional unit (m^2^y). Our life cycle impact assessment revealed that organic farming significantly reduced environmental impacts per tonne of trout in seven out of the nine selected impact categories. Notably, freshwater ecotoxicity exhibited the greatest difference, with organic systems showing a 55% decrease. The only exceptions were freshwater eutrophication and water dependence, where organic production led to higher impacts per tonne of trout. In conventional farming, emissions amounted to 14 kg of P eq./tonne, whereas in organic farming, the emissions were slightly higher (15 kg of P eq./tonne). For water dependence, one tonne of trout production in the conventional system mobilized 128 10^3^ m^3^ vs. 185 10^3^ m^3^ in the organic system. The environmental benefits of organic production were even more marked when using a surface-based functional unit (m^2^y). We demonstrated the benefits of organic trout production from an environmental perspective. However, our findings highlight the caution needed when interpreting LCA comparisons of such production systems that can be highly influenced by methodological choices such as the functional unit used.

## 1. Introduction

Rainbow trout (*Oncorhynchus mykiss*) is the primary farmed fish species reared in France and a significant salmonid species in the global aquaculture production (953,000 tonnes in 2021; FAO, 2022). Only ∼20% of this production is performed in seawater as done in Norway and Chili while the vast majority is coming from freshwater production as practiced in Iran and Turkey, the two main producing countries (FAO, 2023). Traditionally, freshwater trout farming relied on flow-through systems with high water exchange. The lack of space for expansion and new sites (due to competition with other uses and interests), limited freshwater availability, and concerns over the sustainability of the aquafeeds are considered as key obstacles for further expansion of conventional flow-through aquaculture systems (Albrektsen et al., 2022; Chen et al., 2015; Maiolo et al., 2021). As consumer demand for sustainable and environmentally-friendly products grows, there is a rising interest in organic aquaculture, which aims to integrate best environmental practices, natural resource preservation, and high animal welfare standards (Ahmed et al., 2020).

Organic agriculture is often perceived as more sustainable than conventional farming (Meemken and Qaim, 2018). Despite occupying only 1.6% of global agricultural land and accounting for less than 10% of retail sales in most of the countries (Willer et al., 2023), organic farming is one of the fastest-growing sectors in the food industry. Organizations such as the International Federation of Organic Agriculture Movement (IFOAM), the Food and Agriculture Organization (FAO) and the World Health Organization (WHO), through the *Codex Alimentarius*, are working towards establishing an internationally agreed definition of organic practices. In essence, organic farming is an agricultural system that places a high priority on the well-being of ecosystems, encompassing soil, plants, animals, and humans. It relies on ecological processes, biodiversity and cycles adapted to local conditions, rather than the use of inputs with adverse effects. Moreover, organic farming promotes fair relationships and a good quality of life for all involved (IFOAM, 2008). The magnitude of the benefits of organic farming can vary significantly depending on several factors, such as the farm-specific agricultural practices and management approaches, and local environmental conditions (Pépin et al., 2022; Smith et al., 2019). Thus, while organic farming generally fosters environmentally friendly practices, the actual environmental benefits can vary on a case-by-case basis (Meier et al., 2015). Therefore, a comprehensive assessment is necessary to accurately evaluate the overall environmental advantages of organic farming.

Different approaches have been employed to compare the environmental impacts of organic and conventional farming systems, focusing on specific aspects such as biodiversity (e.g., Gabriel et al., 2013; Letourneau and Bothwell, 2008), land use (e.g., Badgley et al., 2007; Connor, 2022; Gibson et al., 2007), or nutrient emissions (e.g., Nowak et al., 2013). However, these assessments offer a limited perspective on the overall environmental impacts of agricultural production. To provide a more comprehensive evaluation, efforts have been made to develop multi-impact methods that can integrate various environmental impact categories, enabling a holistic assessment. The reference method is the Life Cycle Assessment (LCA), which examines the material and energy flows throughout a product’s entire life cycle, encompassing activities like raw material extraction, processing, manufacturing, transportation, distribution, product use, maintenance, recycling, and waste management. LCA is recognized as a comprehensive approach by researchers and international standards (ISO, 2006; Joint Research Centre, 2010) and enables a thorough examination of the different stages and impacts associated with a product’s life cycle.

Tuomisto et al. (2012) and Meier et al. (2015) performed meta-analysis of Life Cycle Assessment (LCA) studies comparing the environmental impacts of organic and conventional terrestrial farming. Their findings indicate that organic farming practices generally yield positive environmental impacts per unit of area, although not necessarily per product unit. Organic production tends to exhibit higher levels of soil organic matter and reduced nutrient losses (such as nitrogen leaching, nitrous oxide emissions, and ammonia emissions). However, when measured per product unit, organic systems were found to have higher levels of nutrient emissions. Additionally, organic systems demonstrated lower energy requirements but higher land use, eutrophication potential, and acidification potential per product unit. Nevertheless, this meta-analysis only concerns land-based production In aquaculture, to the best of our knowledge, only three case studies have been published in peer-reviewed literature: comparisons of conventional and organic production of shrimps (Jonell and Henriksson, 2015) and carp (Biermann and Geist, 2019), and comparison of ingredient types in salmon feeds (Pelletier and Tyedmers, 2007).

This study aims at comparing the environmental impacts of conventional vs. organic rainbow trout farming. To do that, we modelled a trout farm practicing either conventional or organic rearing rainbow trout production. The model we have built aims to simulate a production farm located in France, in Brittany, one of the main rainbow trout producing regions in the country.

## 2. Materials and Methods

### 2.1. Farm model

The farm model, developed using the R freeware (R Development Core Team, 2022), has been partially adapted from previous investigations (Besson et al., 2017, 2016, 2014) to facilitate the acquisition of input values required for conducting a LCA at the farm level. In the present study, the model was customized to simulate, on a daily basis, the production of rainbow trout (*O. mykiss*) in a hypothetical flow-through farm, using actual farm data obtained from surveys conducted in Britany in 2022. The hypothetical farm consisted of 12 concrete raceways, of 100 m^3^ each, for the pre-growing phase, and 24 concrete raceways of 250 m^3^ each for grow-out. Among the 250 m^3^ raceways, 50% received first water, meaning that the water entered the tanks directly from the river, while the remaining 50% received second water, supplied solely by the outlet water from the upstream raceways (Figure 1). In addition to the raceways, the farm was equipped with five feed storage silos and two warehouses measuring 60 and 80 m^2^ (Figure 1). Fish were initially stocked at 10 g and harvested at a fixed weight of 3,000 g that was assumed has the unique market size. The maximal annual production was fixed at 300 tonnes. Throughout the year, three batches of fry were stocked to stagger the sales period (Table 1). We simulated a production over 3 years and used the third year as the reference year for LCA (i.e. year where the first batches stocked in the first year reached market size).

**Figure 1.**
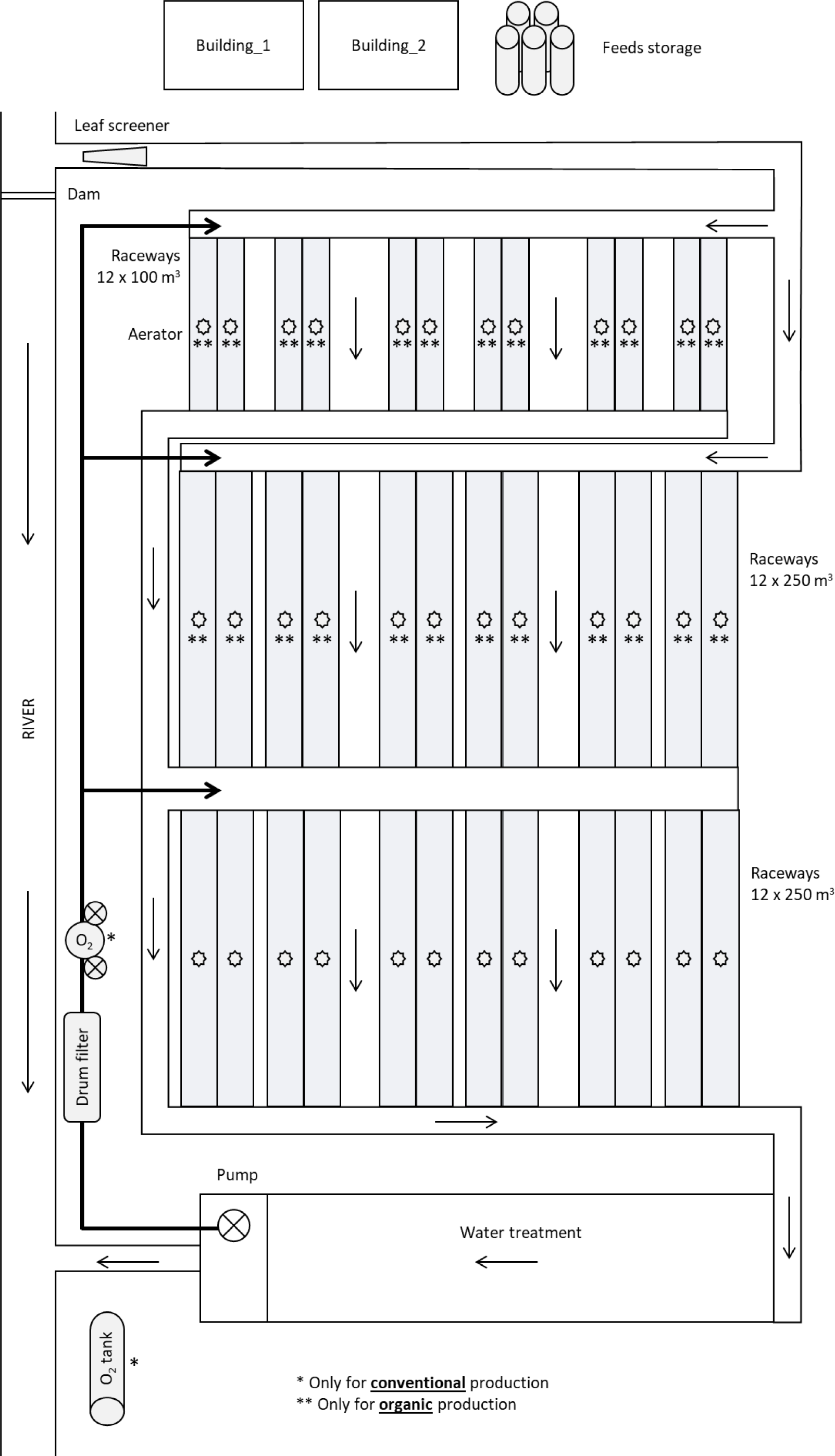
Schematic view of the hypothetical rainbow trout farm used to model conventional and organic production. The equipment specific to the conventional and organic production system are annotated with the following symbols: * and **, respectively.

**Table 1.** Type of trout farms considered in the two different scenarios.

|  | Conventional | Organic |
| --- | --- | --- |
| Production (t year<sup>-1</sup>) | 300 | 203 |
| Rearing duration (d) | 737 ± 2 | 913 ± 4 |
| FCR | 1.3 | 1.3 |
| Mortality rate (%) | 15 | 15 |
| Number of batches per year | 3 | 3 |

FCR = Feed Conversion Ratio calculated as the ratio of feed intake to fish weight gain over one cycle
of production

The various parameters used and the constraints imposed, according to both conventional and organic production scenarios, are elaborated in details below. We incorporated data from surveys, scientific literature, and industry specifications to inform our analysis. Specifically, we used the French production specifications for large trout provided by the Interprofessional Committee for Aquaculture Products (CIPA, 2023) and the regulations for the organic production of aquaculture species established by the French Ministry of Agriculture and Fisheries (MAAP, 2010). A schematic representation of the modelling approach we employed is depicted in Figure 2.

**Figure 2.**
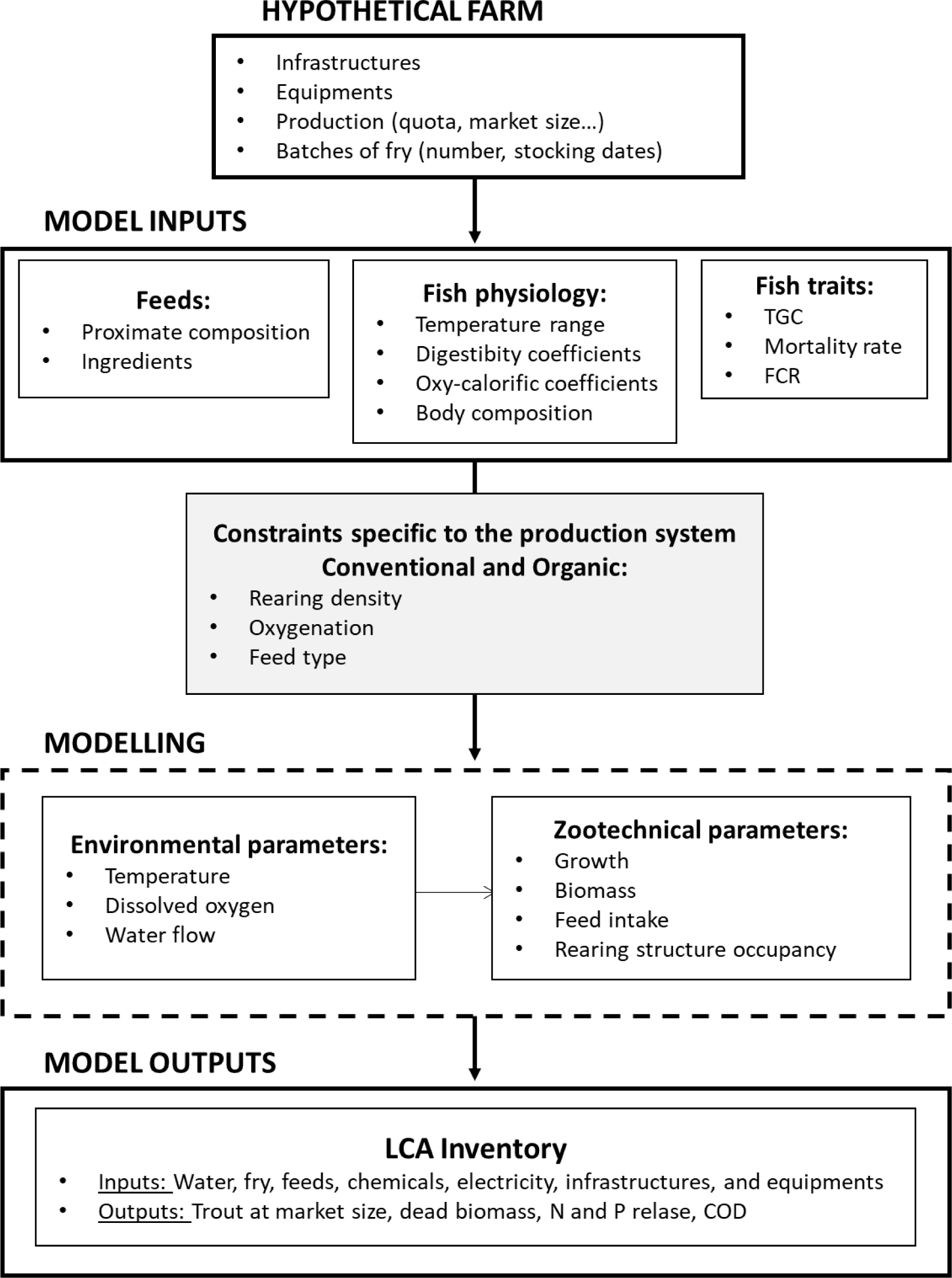
Schematic view of the modelling approach we used. The values for the different model models and details of the constraints applied for the conventional and organic systems are detailed in the text.

#### 2.1.1. Environmental parameters

The daily temperature (T) was modelled using a sinusoidal function with a period of 365 days. As suggested by Seginer and Halachmi (2008), T_n_ is given by:

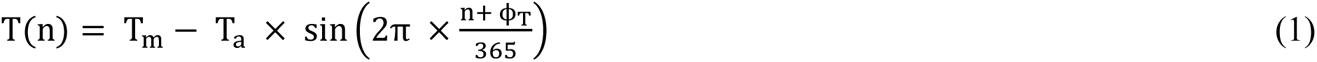

where n is a day from 1 to 365, T_m_ is the mean water temperature (13 °C), T_a_ is the amplitude of the variation (8°C corresponding to a difference of 2 × 8 = 16 °C between the minimum and maximum daily temperature across the whole year) and ϕ_T_ is the phase shift (time-delay of 27.36 d) (Figure 3A).

**Figure 3.**
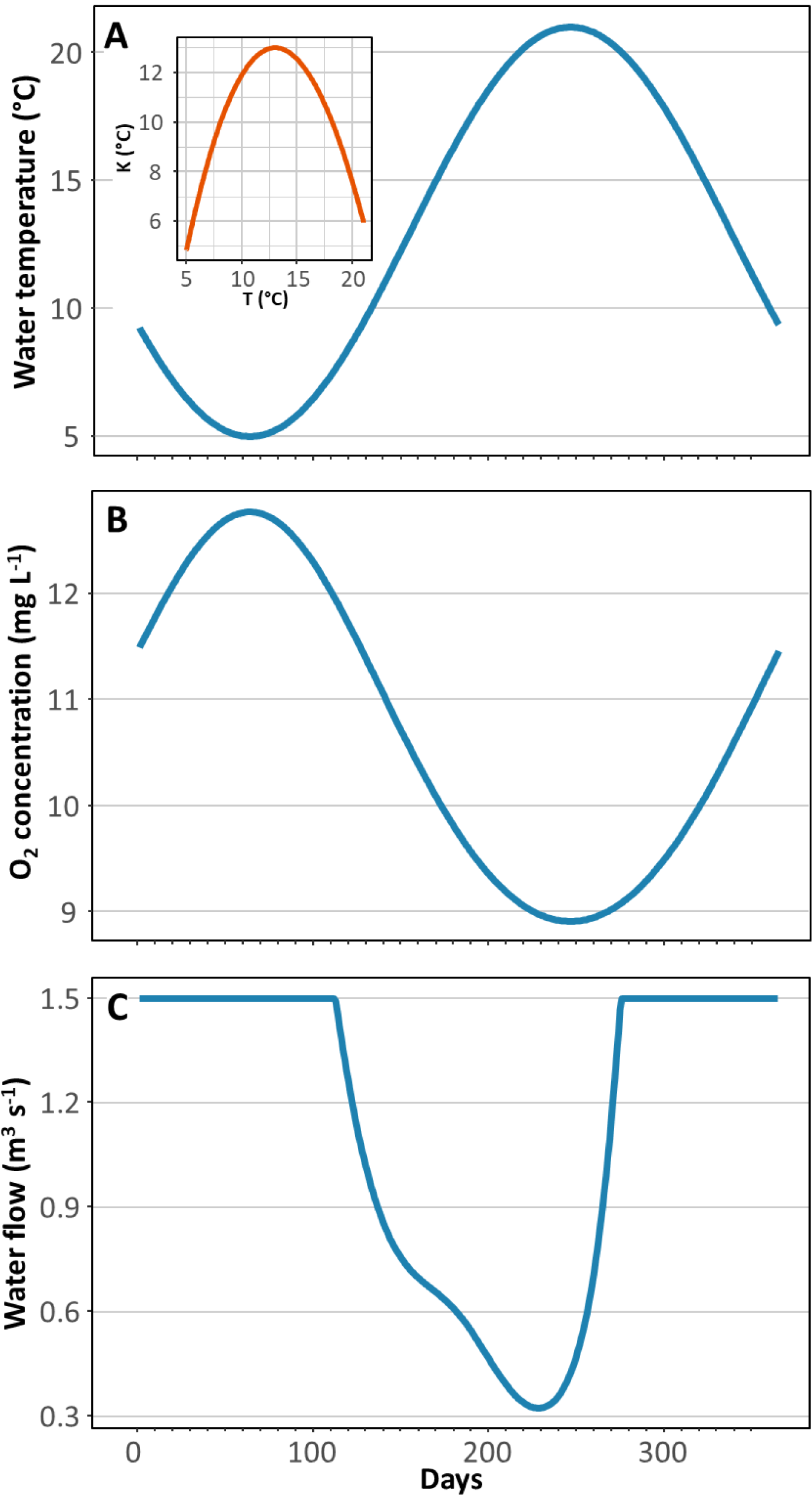
Graphical representations of (A) the simulated annual temperature conditions with, in insert, corrected temperature K as a function of temperature T and (B) the resulting oxygen concentration in water. (C) the simulated annual water flow entering the fish farm.

Dissolved oxygen concentration ([O_2_] in mg L^-1^; Figure 3B) at day n in surface water was calculated from Mortimer (1956) considering a standard pressure of 1 atm:

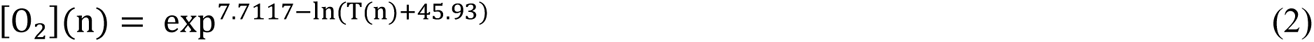

where T_n_ is the daily temperature (in °C).

The water flow within the fish farm, which experiences fluctuations throughout the year, was simulated based on actual water flow data obtained from a river in Brittany. Data from the years 2018 to 2022, specifically from the Aulne River in Brittany, were collected from the reference HydroPortail database version 3.1.4.3 (HydroPortail, 2023). Two constraints were considered when calculating the water flows: the inflow into the fish farm could not exceed 1.5 m^3^ s^-1^, and a maximum of 90% of the total river flow could be derived to the fish farm. To predict the daily water inflows into the fish farm, a Generalized Additive Model (GAM) was then employed considering the different constraints (Figure 3C).

#### 2.1.2. Growth

The fish model described the daily weight and the daily weight gain of fish based on thermal growth coefficient (TGC). Considering that the relationship between growth rate and water temperature is non-linear, the TGC formula was corrected for the concave relationship between growth rate and temperature, using a corrected temperature K (Mallet et al., 1999) as suggested by Besson et al. (2016):

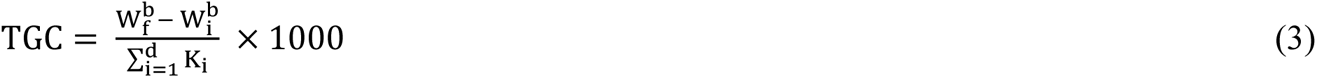

where W_f_ represents the final weight at harvest (3000 g), W_i_ denotes the initial weight at stocking (10 g), d is the rearing time in days and b is a weight coefficient set at 1/3 for the overall growing period even if this parameter can vary according to growth (Dumas et al., 2007).

The TGC values were adjusted to 1.80 and 1.45 (g^1/3^ °C^-1^ d^-1^) in the conventional and organic production scenarios, respectively. We simulated a 24-month production cycle in conventional production and a 30-month production cycle in organic production (Figure 4). This rearing time difference corresponds to the expected growth differentials between triploid monosex trout, primarily used in conventional production, and male and female diploid trout (Aqualande Origins, 2019) used in organic production according to regulatory requirements (MAAP, 2010). In the conventional production scenario, the storage dates were kept constant throughout the three years and set at d 30 for the first batch, followed by intervals of 100 days (i.e., d 130 for batch 2 and d 230 for batch 3) over the course of a year. In the organic production scenario, the frequency of batch entries was set at 50 days (i.e., d 80 for batch 2 and d 130 for batch 3) to maintain the same rotation of harvests and stocking (3 entries and 3 harvests per year). This adjustment was necessary to accommodate the longer rearing duration (i.e. 30 vs 24 months) while ensuring consistent batch rotation in the organic production system.

**Figure 4.**
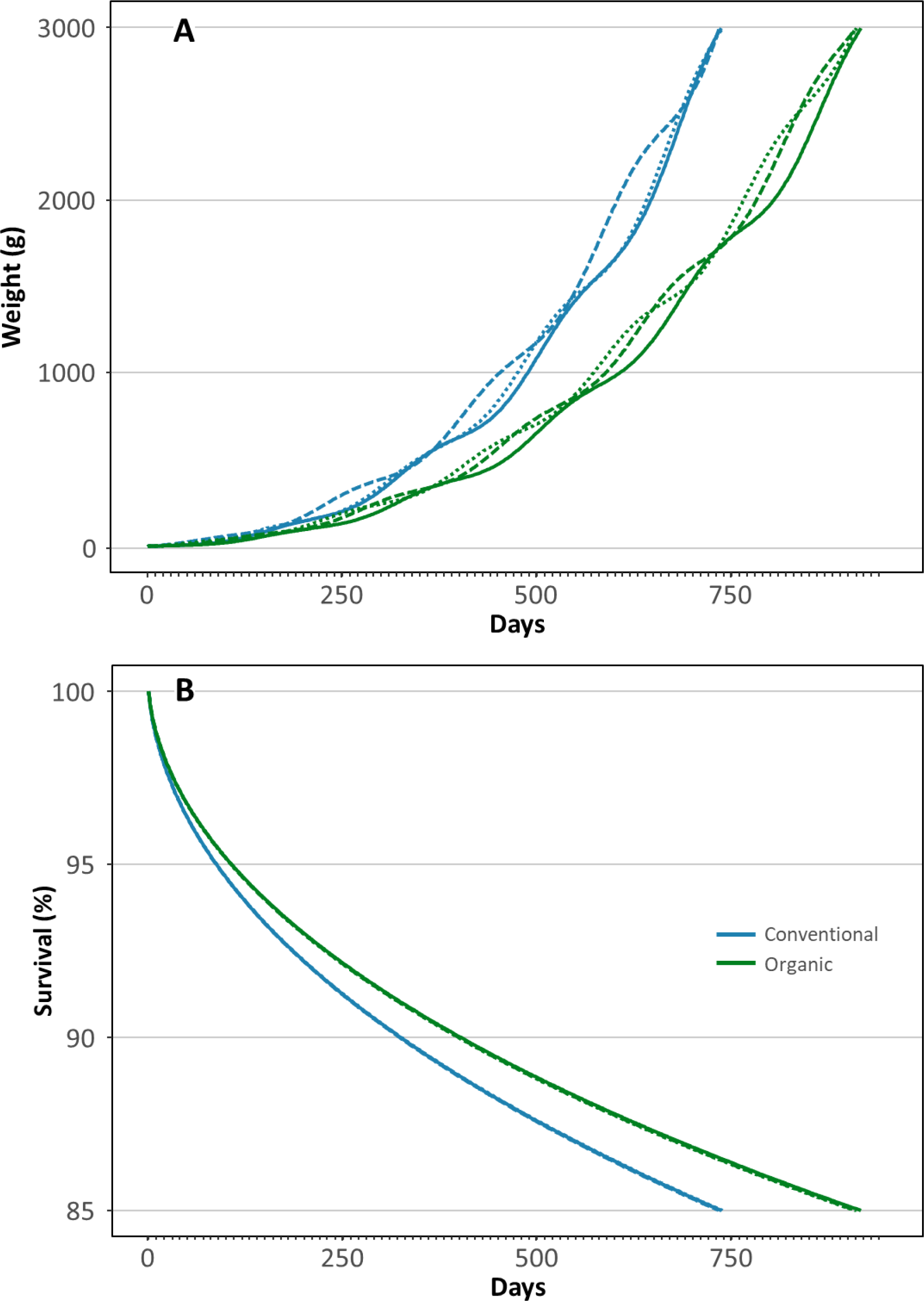
Graphical representations of (A) growth performances, from 0.01 to 3 kg and (B) survival of the three fish batches in conventional and organic production systems.

The corrected temperature (K) at a given day n was calculated as follows:

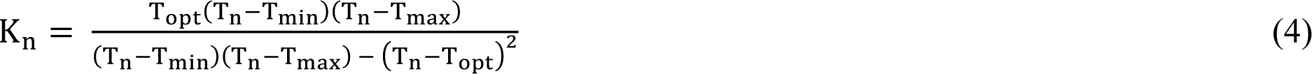

where T_min_ ≤ K ≤ T_max_ and K = 0 for other values. Here, T_min_ and T_max_ represent the minimum and maximum temperatures, respectively, below and above which growth does not occur. T_opt_ refers to the optimal temperature for growth. Based on extrapolations from Bear et al. (2007), the values for rainbow trout were set at 3 °C for T_min_ (K = 0), 13 °C for T_opt_ (K = 13), and 24 °C for T_max_ (K = 0). Consequently, for a positive growth rate, T_n_ must fall between 3 °C and 24 °C. The daily weight (W) and daily weight gain (DWG; g d^-1^) can be calculated as follows at day n:

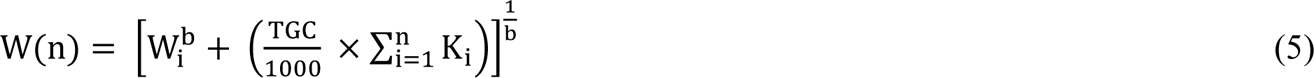

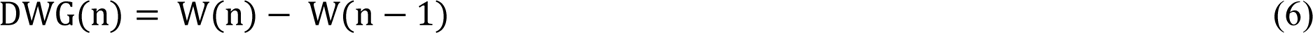

Growth curves under the two production scenarios are presented in Figure 4A.

#### 2.1.3. Mortality

In this study, a mortality rate of 15% was applied throughout the entire production cycle, spanning from 10 to 3000 g. It was assumed that the probability of daily mortality was not linear across the rearing period and is higher for younger individuals (Gåsnes et al., 2021). To model this, a Weibull function was considered for the lifetime distribution, as it is commonly used for survival analysis (Carroll, 2003). So, the hazard function h which defines the death rate at a given day (n) conditional on survival until time n or later can be calculated as follows:

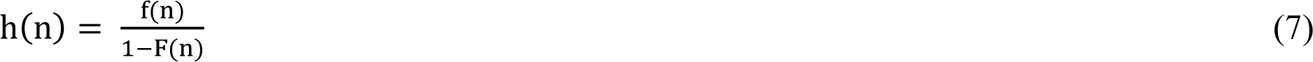

considering the Weibull density function

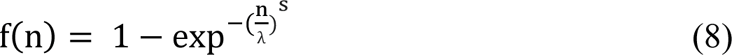

and the Weibull distribution function

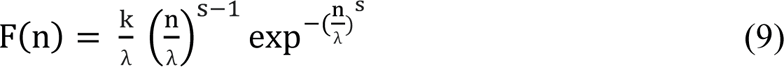

While the shape parameter (s) was kept fixed at 0.5, the scale parameter (λ) was optimized for each fish batch, ensuring a final mortality rate of 15% across the entire rearing duration.

#### 2.1.4. Biomass

The biomass (BM) at a given day for each batch was determined as follows:

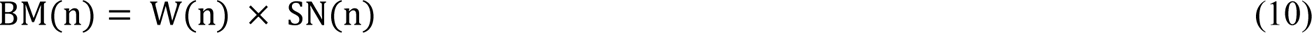

where W is the individual body weight at a given day n and SN the number of surviving fish at this day. In the same way, the dead biomass at day n was calculating by replacing SN by the number of dead fish at this day in the equation (10).

The total production (in tonnes) was then calculated as the difference between the biomass at the harvest and the initial biomass at stocking. In the two different production scenarios, harvest took place at a constant weight of 3000 g.

#### 2.1.5. Raceways occupation

In our model, the occupancy of the raceways was determined by the densities achieved, which necessitates regular sorting of the fish during rearing. Initially, we assumed that each batch was stocked in a 100-m^3^ raceway. As the fish grow, they were periodically redistributed into 2 and then 4 raceways of 100 m^3^ before ultimately occupying 4 then 8 raceways of 250 m^3^. The maximum density constraints varied depending on the production scenario. In the conventional production scenario, the density limits applied were 50 kg m^-3^ when W ≤ 50 g, 70 kg m^-3^ when 50 g < W ≤ 1000 g and then 90 kg m^-3^ when W > 1000 g (CIPA, 2023). For the organic production scenario, the density limits were as follows according to CIPA (2023): 25 kg m^-3^ when W ≤ 15 g, 30 kg m^-3^ when 15 g < W ≤ 30 g and then 35 kg m^-3^ when W > 30 g. The percentage of occupancy of each rearing structure was calculated as the sum of the surface used per day divided by the total surface available over a year (expressed as m^2^y).

#### 2.1.6. Feeds

Feed conversion ratio at a given day (FCR) was modelled by a third-order polynomial model based on fish body weight (W) using equation extrapolated from Bureau and Hua (2008) :

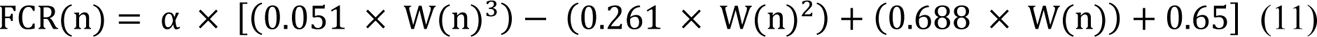

where α is a scaling factor to obtain a realized FCR of 1.30 kg kg^-1^ over the production cycle for each batch in the two production scenarios assuming that the conventional and organic fish lines have the same feed efficiency (Figure 5). Daily feed intake (DFI, kg d^-1^) is calculated back from FCR and DWG by:

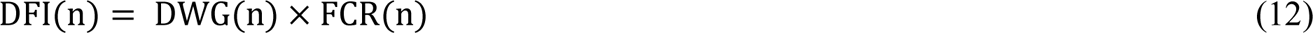

**Figure 5.**
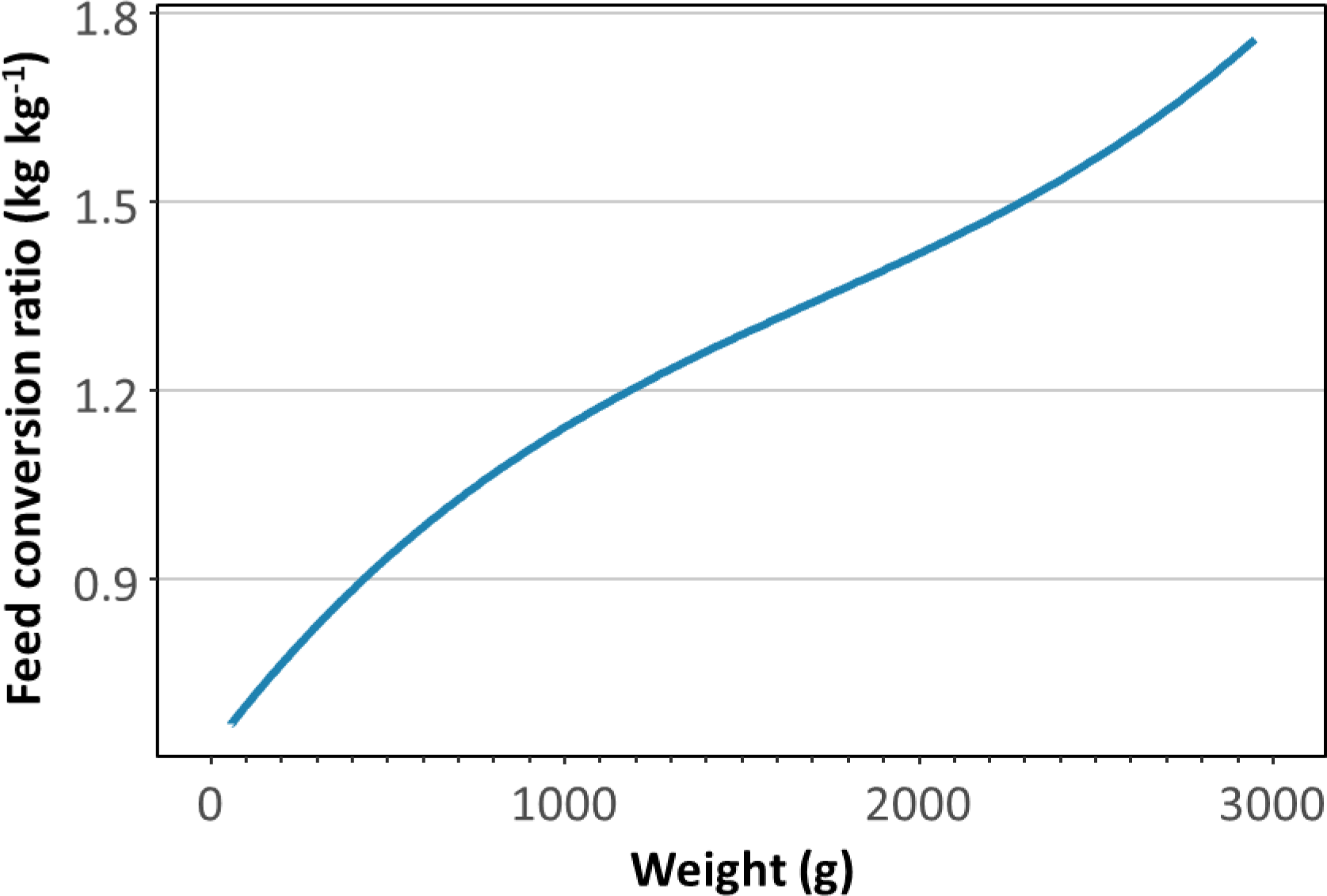
Estimated FCR of rainbow trout at increasing live weight extrapolated from Bureau and Hua (2008).

In our model, we considered the dynamic nature of fish feed composition, particularly in terms of protein and lipid content, throughout the rearing period. As a result, four different types of feed were incorporated based on the weight. Feed 1 was used for fish up to 50 g, feed 2 for fish up to 500 g, feed 3 for fish up 1500 g, and finally, feed 4 was used until reaching the harvest weight (W_f_). This approach ensures that the nutritional needs of the fish are adequately met at each stage of their growth and development. Conventional or certified organic feeds were used depending on the production scenario.

#### 2.1.7. Nutrient release

The concentration of nutrients (N and P) and chemical oxygen demand (COD) in effluent water was determined using a mass-balance approach (Aubin et al., 2011). To model excretion, the first step involved calculating the total nutrient amount provided by the feeds (N_feed_), taking into account two fractions: the portion consumed (N_eaten_) and the portion wasted (N_waste_) on day n, along with the nutrient fixation by the fish (N_fish_). It was assumed that 1% of the distributed feeds remained uneaten (Boujard et al., 1995). The proximate composition of the different feeds can be found in Table 2.

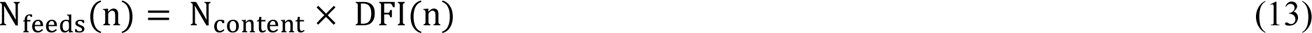

**Table 2.** Composition of the feeds used in the two different scenarios.

|  | Conventional | Organic |
| --- | --- | --- |
| Proteins (%) |  |  |
| Feed 1 | 45 | 43 |
| Feed 2 | 40 | 39 |
| Feed 3 | 39 | 38 |
| Feed 4 | 38 | 36 |
| Lipids (%) |  |  |
| Feed 1 | 21 | 21 |
| Feed 2 | 23 | 24 |
| Feed 3 | 27 | 26 |
| Feed 4 | 30 | 28 |
| Carbohydrates (%) |  |  |
| Feed 1 | 12.0 | 13.0 |
| Feed 2 | 13.9 | 14.0 |
| Feed 3 | 12.8 | 13.6 |
| Feed 4 | 12.8 | 11.4 |
| Phosphorus (%) |  |  |
| Feed 1 | 0.95 | 1.70 |
| Feed 2 | 0.95 | 1.70 |
| Feed 3 | 0.90 | 1.70 |
| Feed 4 | 0.90 | 1.60 |

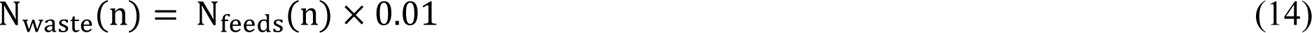

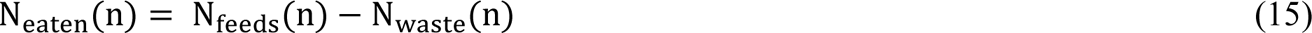

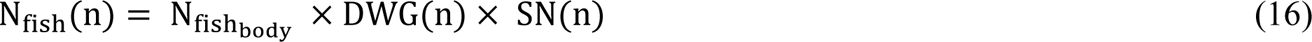

where N_fish_body__ is the nutrient composition of the fish (in kg kg^-1^) set at 0.03 for N (Oz and Dikel, 2015) and 0.004 for P (Kause et al., 2022).

The total nutrient excretion (N_excretion_) was given by:

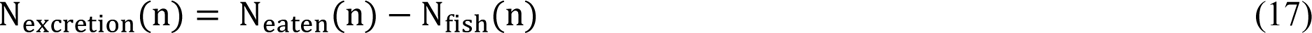

Calculation of the suspended (N_suspended_) and dissolved (N_dissolved_) was given by:

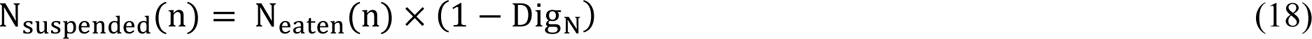

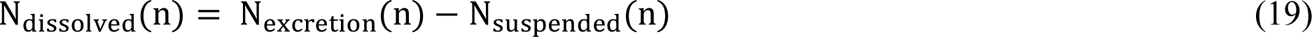

where Dig_N_ is the digestibility coefficient set at 94% for proteins and 61% for P (Dalsgaard and Pedersen, 2011).

The final amount of N release was then calculated considering that the sedimentation area used as water treatment is able to remove 20% of suspended N (Stewart et al., 2006):

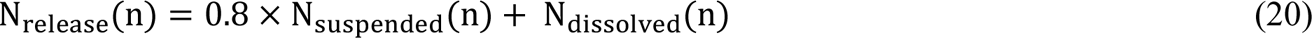

COD at a given day n was calculated using feed quantity eaten (DFI) at day n, the proximate protein, lipids and carbohydrates contents of the feeds (P_feeds_, L_feeds_ and C_feeds_) and their respective digestibility (Dig) (i.e., 94% for proteins, 91% for lipids and 67% of carbohydrates; Dalsgaard and Pedersen, 2011):

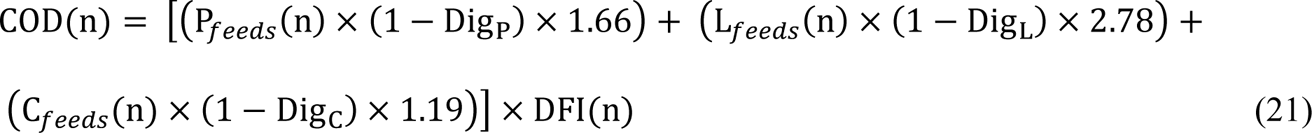

where the coefficients applied for protein, lipids and carbohydrates were coming from Meriac et al. (2014).

#### 2.1.8. Oxygen

In both production scenarios, the primary constraint for oxygen management was to maintain a saturation level of 80% at the outlet of the raceways. However, the approach to O_2_ supplementation differed between the two production scenarios. In conventional production, liquid oxygen was used for O_2_ supplementation, whereas in organic production, the use of aerators was the only permissible method (MAAP, 2010). In our model, the amount of oxygen added was determined based on the difference between the supply of oxygen through the water inlet (O_2inlet_), which could come directly from the river or from the upstream raceways (Figure 1), and the O_2_ consumption by the fish (O_2cons_). These two parameters were calculated using the following equations:

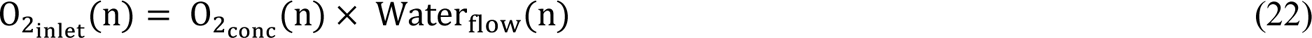

where O_2conc_ is the O_2_ concentration from water inlet either coming from the river - in this case O_2conc_ = [O_2_](n) (see Section 2.1.1) or from the upstream raceway – in this case O_2conc_ = [O_2_](n) - O_2cons_(n) of the upstream raceway. Water_flow_ in a given raceway the water flow through the raceway calculated as follows:

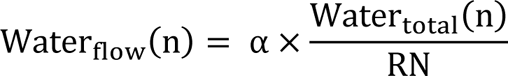

where Water_total_ is the water flow for the whole fish farm, RN is the number of raceways and α is a size coefficient (i.e. 0.29 for 100-m^3^ raceway and 0.71 for 250-m^3^ raceway). Then, O_2_ consumption is given by:

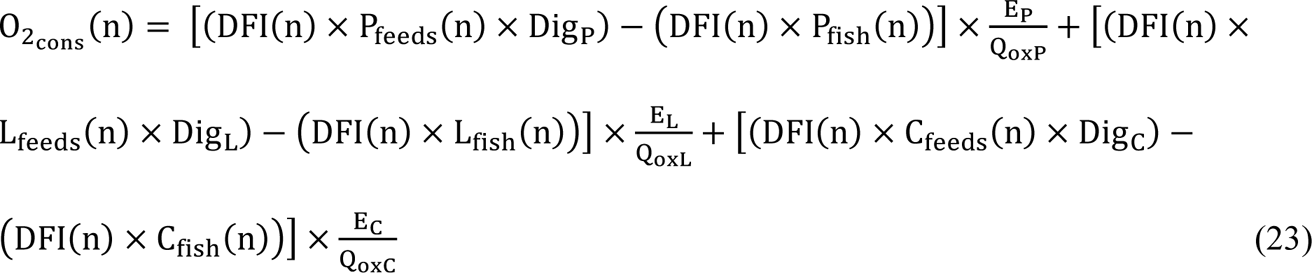

where Q_oxP_, Q_oxL_ and Q_oxC_ are the oxy-caloric coefficients of proteins (13.4 MJ kg O_2_ ^-1^), lipids (13.7 MJ kg O_2_ ^-1^) and carbohydrates (14.8 MJ kg O_2_ ^-1^) (Brafield and Solomon, 1972; Elliott and Davison, 1975) and E_P_, E_L_ and E_C_ are the energy contents of proteins (23.6 MJ kg^-1^), lipids (39.5 MJ kg^-1^) and carbohydrates (17.2 MJ kg^-1^) (Brafield and Llewellyn, 1982).

If the difference between O_2inlet_ and O_2cons_was higher than the 80% saturation O_2_ concentration (O_2_80%_ = 0.8 [O_2_]), it indicated that no oxygenation or aeration is required. Conversely, a result lower than O_2_80%_ indicated the need for O_2_ supplementation (O_2sup_) either through the addition of liquid O_2_ or by aeration:

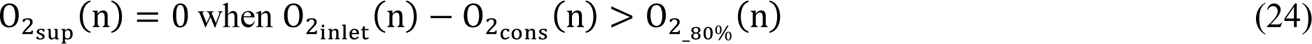

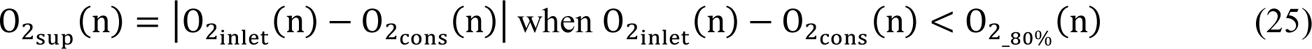

#### 2.1.9. Energy

The electricity consumption of the farm was modelled taking into account water filtration, oxygenation and recirculation processes. A drum filter (1 kWh) and a recirculation pump (20kWh) operated during periods when the water flow was at its lowest, typically between May and September. Their purpose was to ensure effective water recirculation during this period under both conventional and organic production scenarios. Electricity consumption by the filter (E_filter_) and the recirculation pump (E_pump_) at a given day n has been calculated as follows:

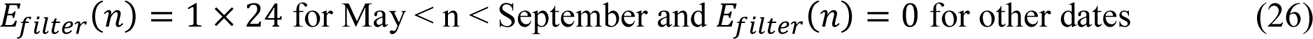

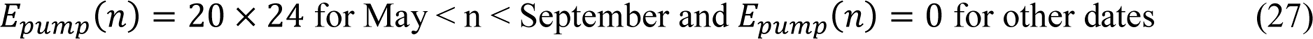

One key distinction between estimating electricity consumption for conventional and organic production lies in the method employed for water oxygenation. In conventional production, liquid oxygen was added using an oxygen cone and two pumps with a power consumption of 20 kWh each. In this case, the electrical consumption at a given day n was calculated as follows:

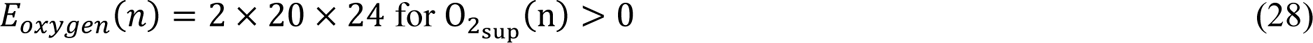

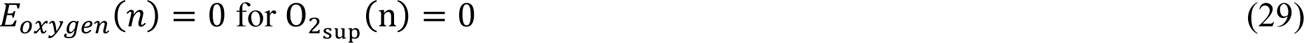

In the organic production scenario, the use of aerators replaced liquid oxygen. These aerators enable the addition of 1.5 kg of oxygen per kilowatt-hour (kWh) of electricity consumed (Ahmad and Boyd, 1988; Brown et al., 2014). Consequently, the calculation for electrical consumption associated with the aerators has been calculated as follows:

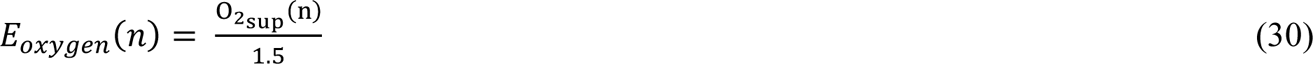

The total electricity consumption (E_total_) was determined by summing the electricity usage for water filtration, oxygenation, and recirculation:

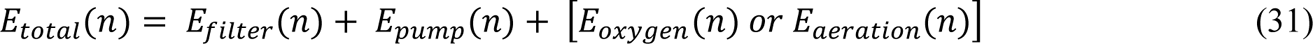

### 2.2. Life Cycle Assessment (LCA)

#### 2.2.1. Goal and scope

An attributional LCA was conducted according to the general requirements of the methodology proposed by ILCD standards (Joint Research Centre, 2010). The methodology was adapted to the characteristics of fish farming. The goal and scope of this study was the environmental assessment of trout farming in a hypothetical farm producing large rainbow trout following either (1) conventional or (2) organic practices in the same infrastructures. The system was defined from cradle-to-farm-gate and included five distinct sub-systems (Figure 6): (1) production of purchased feed, including cultivation of ingredients, processing, and transportation; (2) production of energy expended at farm level (electricity); (3) production of farming facilities and equipment used; (4) chemicals, including liquid oxygen, veterinary and disinfection products, and their transportation (5) farm operations, including nutrient emissions from the biological transformation of feed after onsite treatment of wastewater (see details in Section 2.2.2). The functional unit in which environmental impacts were expressed was one tonne of trout produced at farm level on a basis of one year of routine production. We also expressed the environmental impacts using an surface-based functional unit (m^2^y) as recommended by Van der Werf et al. (2020). Here we considered only the surface directly involved in the fish production.

**Figure 6.**
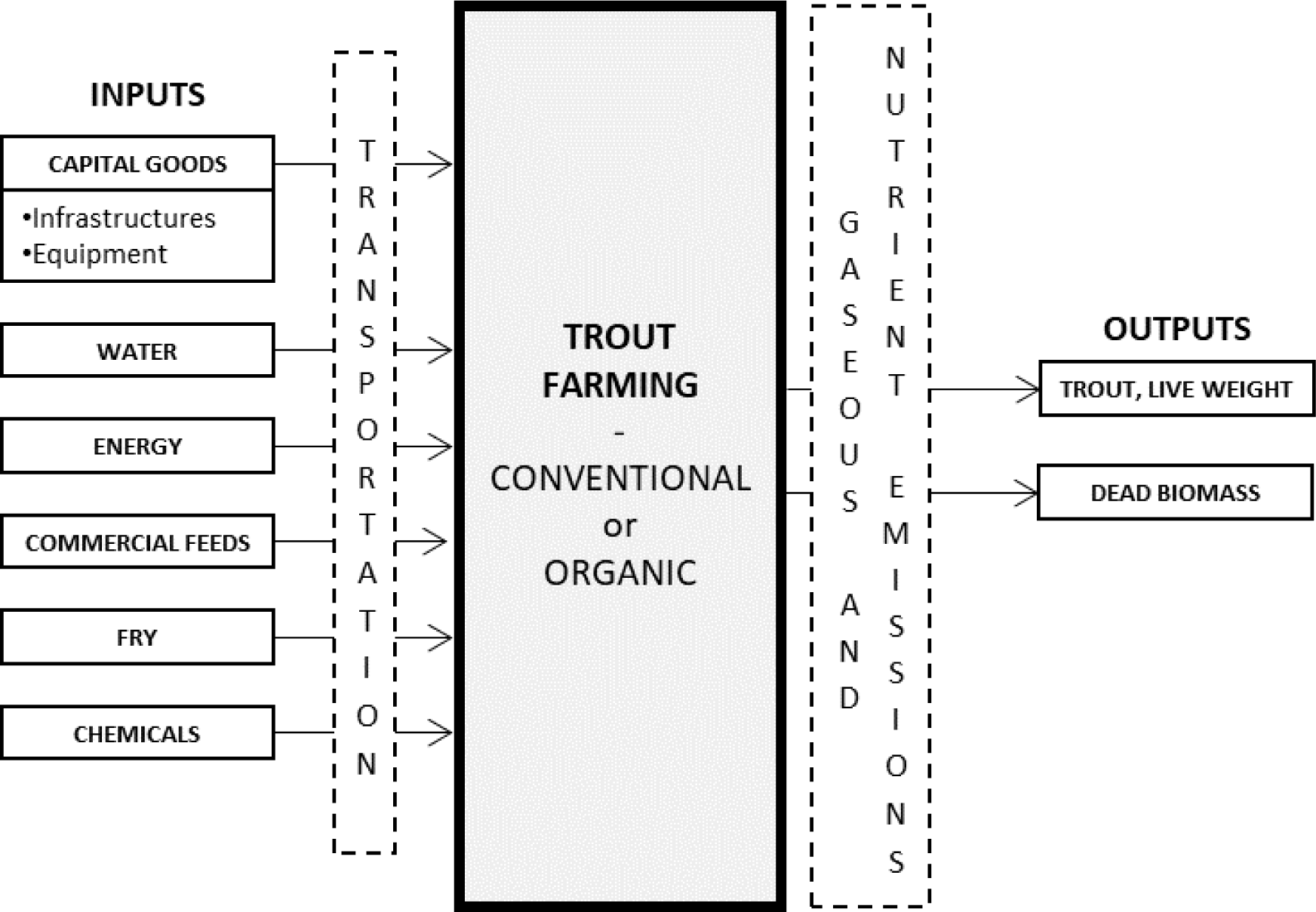
System boundaries and flows of rainbow trout *Oncorhynchus mykiss* grow-out production.

#### 2.2.2. Life cycle inventory

The life cycle inventory (LCI), presented in Table 3, was conducted by running our farm model with the specifications for both conventional and organic production scenarios. All the inputs and outputs were calculated using all the results from each batch of fish over one year of routine production generated as described in the farm model. The Agribalyse version 3.0 (Koch and Salou, 2022) and Ecoinvent version 3.8 (Wernet et al., 2016) databases were used to obtain the necessary data for conducting the assessment. Both databases are grounded on the recommendations in international standards (Wolf et al., 2012).

##### (1) Production of purchased feed

Crop-derived ingredients used in fish feed mainly originated from Brazil and France (e.g. soybean meal from Brazil and wheat bran from France). Fish-derived ingredients originated from the Peruvian and the Norwegian fish milling industry (e.g. fish meal from Peru and fish meal from fish trimming from Norway). The exact composition of the different feeds used and their nutritional values were given by the feed manufacturer (Le Gouessant, personal communication). The transport of feed ingredients to feed manufacturers in France was by trans-oceanic ship and by lorry (>32 t), whereas the transport of feed from France to the fish farm in Brittany was by lorry (>32 t). Road distances were calculated from Google Maps and ocean distances were assessed from shiptraffic.net. Other data required to compute the environmental impact of feed ingredients were based on the literature (Boissy et al., 2011; Pelletier et al., 2009).

##### (2) Production of energy expended on the farm

The electricity used by the farm was coming from the French energy mix in the Ecoinvent database. Annual on-site consumption from other energy sources (diesel and gas) where considered negligible.

##### (3) Production of farming facilities and equipment used

We considered the construction of two different buildings with a life span of 30 years. Nevertheless, the life span of each rearing structures has been adjusted in LCA inventory according to the rearing structures’ occupancy (Table 4) calculated as described in Section 2.1.5 assuming that the actual life span of the rearing span is related to their level of occupancy. The production of equipment used (i.e. pump, tanks) was calculated using data from INRAE.

##### (4) Chemicals

This sub-system included the veterinary and disinfection products. While the use of these products varies little between conventional and organic production, the main difference is the inclusion in this sub-system of the liquid oxygen used only in conventional production. Here we considered production of liquid oxygen from cryogenic air separation process.

##### (5) Farm operations

The farm operation sub-system included the use of facilities and equipment and the emissions of pollutants from the biological transformation of the feed distributed to the fish. The amount of nitrogen (N), phosphorus (P) and chemical oxygen demand (COD) of the dissolved organic matter excreted by the fish in effluent water were calculated through mass balance (Papatryphon et al., 2005) considering the onsite treatment capacity of the sludge settling pond. Sludge produced by the farm was used for neighbourhood agricultural purposes and was not included in the analysis.

**Table 3.** Life Cycle Inventory for one year of production.

|  | Item | Unit | Conventional | Organic |
| --- | --- | --- | --- | --- |
| INPUTS | Site surface | m<sup>2</sup> | 16000 | 16000 |
|  | Water | m<sup>3</sup> | 35785586 | 35785586 |
|  | Fry (10 g) |  |  |  |
|  | Triploid trout (♀) | u | 120000 | - |
|  | Organic trout (♀/♀) | u | - | 81000 |
|  | Feeds |  |  |  |
|  | feed_1/ feed_org_1 | kg | 3127 | 2120 |
|  | feed_2/ feed_org_2 | kg | 41192 | 29841 |
|  | feed_3/ feed_org_3 | kg | 120688 | 69479 |
|  | feed_4/ feed_org_4 | kg | 223868 | 151370 |
|  | Chemicals |  |  |  |
|  | Liquid oxygen | m<sup>3</sup> | 277036 | - |
|  | Antibiotics | kg | 0.24 | 0.16 |
|  | Others | kg | 4000 | 4000 |
|  | Electricity | kWh | 427512 | 106440 |
|  | Infrastructures |  |  |  |
|  | 60-m<sup>2</sup> building | u | 1 | 1 |
|  | 80-m<sup>2</sup> building | u | 1 | 1 |
|  | 100-m<sup>3</sup> raceways | u | 12 | 12 |
|  | 250-m<sup>3</sup> raceways | u | 24 | 24 |
|  | Equipment |  |  |  |
|  | Feed storage silo | u | 5 | 5 |
|  | Oxygen cone | u | 2 | - |
|  | Oxygen tank | u | 1 | - |
|  | Leaf screener | u | 1 | 1 |
|  | Fish elevator | u | 2 | 2 |
|  | Drum filter | u | 1 | 1 |
|  | Electric generator | u | 1 | 1 |
|  | Pumps | u | 3 | 1 |
|  | Aerators | u | 12 | 36 |
|  | PVC pipe | m | 1500 | 1500 |
| OUTPUTS | Trout at market size (3 kg) | kg | 300478 | 202909 |
|  | Dead biomass (incinerated) | kg | 9158 | 5990 |
|  | Water (back to river) | m<sup>3</sup> | 35785586 | 35785586 |
|  | Nitrogen (in river) | kg | 14512 | 8793 |
|  | Phosphorus (in river) | kg | 2254 | 2701 |
|  | COD (in river) | kg | 62435 | 39631 |

u: unit; COD: Chemical Oxygen Demand; Transportation was included at each step when needed.

**Table 4.**
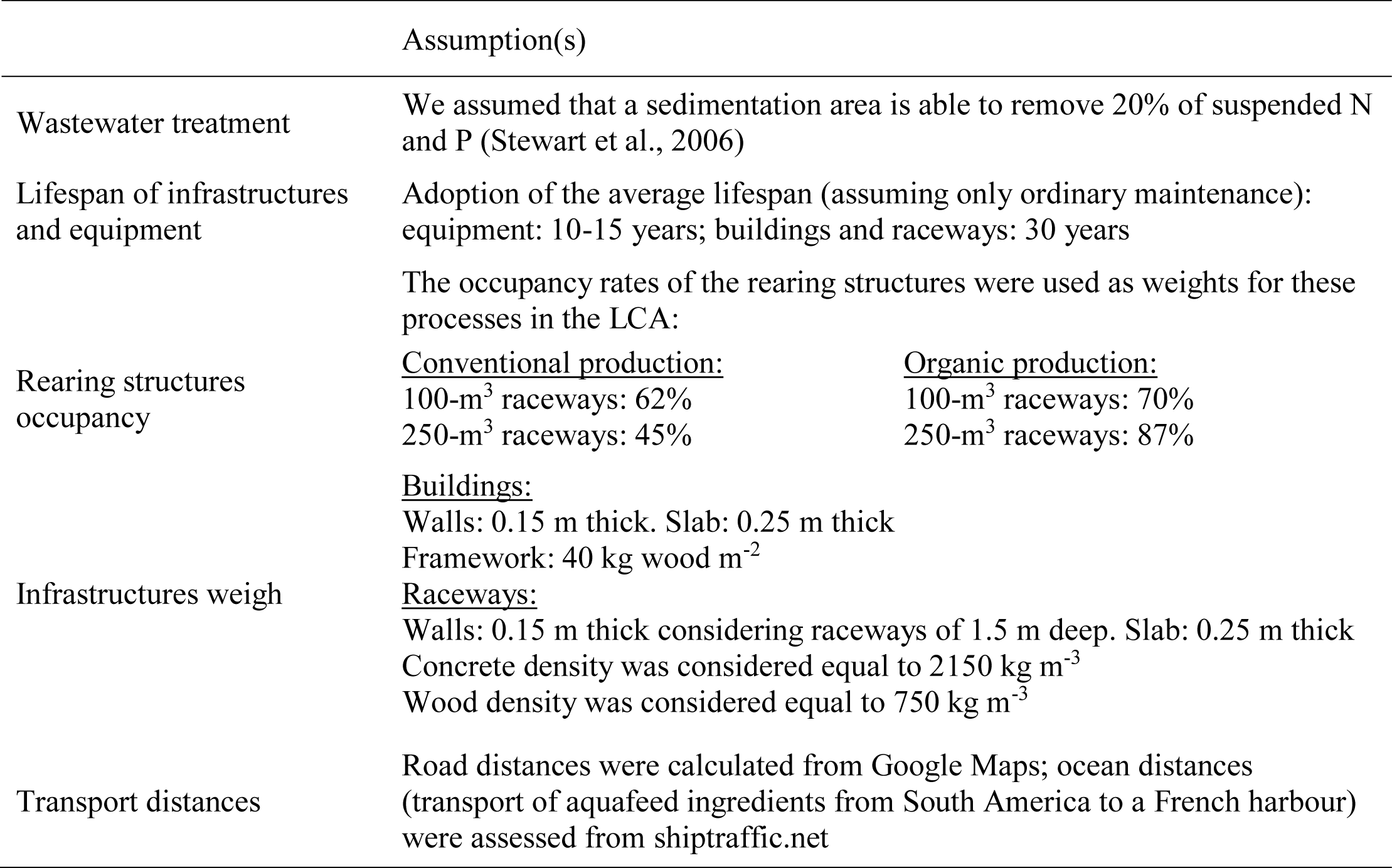
Assumptions made to fill inventory gaps.

Gaps in the inventory were filled on the basis of the assumptions reported in Table 4.

#### 2.2.3. Life cycle impact assessment

The assessment of the impact was carried out using ReCiPe 2016 Midpoint (H) version 1.07 (Huijbregts et al., 2017), which is a methodology based on the Eco-indicator and CML approaches. According to the European Commission/JRC (2010), ReCiPe represents the most up-to-date and standardized indicator approach available for life cycle impact assessment.

Table 5 provides a breakdown of the nine selected impact categories from ReCiPe, namely climate change (GWP), terrestrial acidification (TAP), freshwater eutrophication (FEP), marine eutrophication (MEP), terrestrial ecotoxicity (TETP), freshwater ecotoxicity (FETP), land use (LU), water dependence (WD) and the Cumulative Energy Demand method (CED; Frischknecht et al., 2007). These impact categories have been identified among the most suitable indicators of aquaculture impacts (Bohnes et al., 2019). To enable comparison with previous studies on trout production systems, the CML baseline (Guinée, 2002) was used as an alternative to the ReCiPe approach. The environmental impacts were calculated using Simapro version 8.0 software (PRé Consultants, 2014).

**Table 5.** Characteristics of the selected impact categories.

| Impact category | Abbreviation | Unit | Definition |
| --- | --- | --- | --- |
| Climate change potential | GWP | kg CO<sub>2</sub> eq. to air | the contribution of greenhouse gases to global warming |
| Terrestrial acidification potential | TAP | kg SO<sub>2</sub> eq. to air kg | changes in acidity in the soil due to a change in acid deposition, which in turn is a consequence of changes in air emission of NO<sub>x</sub>, NH<sub>3</sub> and SO<sub>2</sub> |
| Freshwater eutrophication potential | FEP | kg P eq. to freshwater | a change in the levels of P in freshwater caused by emissions of nutrients into water and soil |
| Marine eutrophication potential | MEP | kg N eq. to freshwater | a change in the levels of N in marine water caused by emissions of nutrients into water and soil |
| Terrestrial ecotoxicity potential | TETP | kg 1,4-DCB eq. to soil | a change in the levels of toxic chemicals caused by emissions into the soil |
| Freshwater ecotoxicity potential | FETP | kg 1,4-DCB eq. to freshwater | a change in the levels of toxic chemicals caused by emissions into the water |
| Cumulative energy demand | CED | GJ eq. | the direct and indirect consumption of energy |
| Land use | LaU | m<sup>2</sup>y crop eq. | the ground surface used directly (land occupied by ponds) and indirectly (land used to grow feed ingredients) |
| Water dependence | WD | m<sup>3</sup> | the water flowing into the production system |

#### 2.2.4. Sensitivity analysis

Considering that the feed use is the major contributors to environmental impacts, a sensitivity analysis was conducted on FCR. In this study, we ran the model, both for the conventional and organic productions, to gauge the changes in the different LCA impact categories when FCR varied from 1.0 to 1.6 in steps of 0.1.

## 3. Results

Figure 7 presents the level of the environmental impacts and the contribution of the system components to the impacts for conventional and organic productions of rainbow trout. The impacts are calculated according to ReCiPe method using two different functional units: per tonne of trout (product-based) and per m^2^y (surface-based). Among the nine impact categories analyzed, the conventional production system exhibits higher impacts for all categories, except for FEP and WD when the results are expressed per tonne of trout. For instance, in the conventional production system, one tonne of trout emits 14 kg P eq. and depends on 128,000 m^3^ of water, while an equivalent quantity of organic trout emits 19 kg P eq. and depends on 185,000 m^3^ of water. Obviously, when the results are expressed per m^2^y, organic production shows lower environmental impacts for all considered categories, including FEP and WD (Figure 7). Regardless of the functional units chosen (product-based or surface-based), other impact categories also follow similar trends, with the surface-based functional unit leading to a larger gap between the two production scenarios. The results presented in Figure 7 cannot be compared with those presented in previous LCA publications, which were mostly obtained with the CML baseline method. Thus, the environmental impacts of the production of one trout were also assessed using the CML baseline methodology and compared with literature for conventional production (Table 6). Overall, our results are consistent with those found in previous studies. For the sake of clarity, the values presented in the subsequent paragraphs are given for the ReCiPe method and per tonne of trout at market size, while the results expressed per m^2^y can be found in Figure 7.

**Figure 7.**
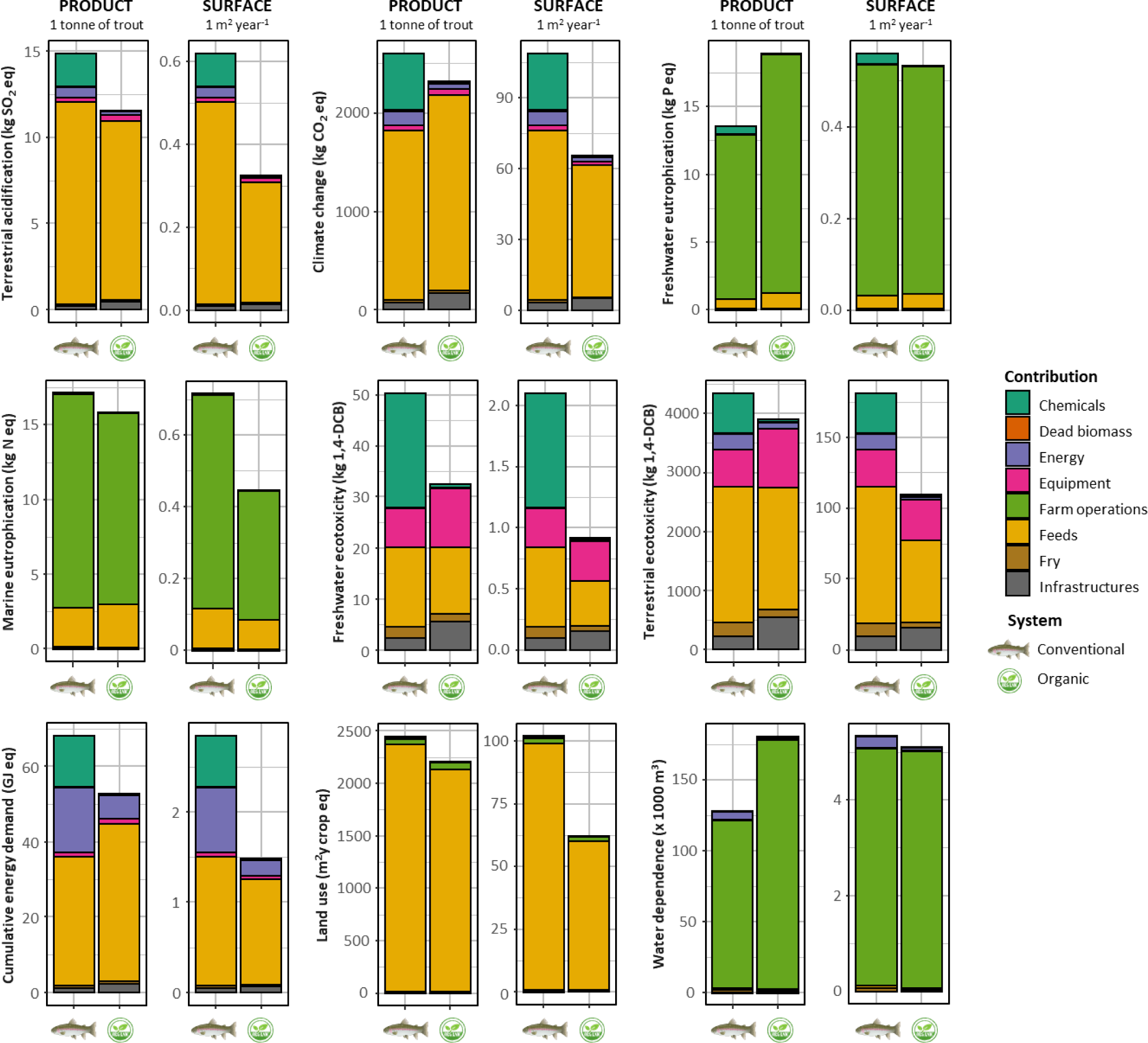
Contribution of each input or production step in environmental impacts in conventional and organic fish production system. Results are either expressed per tonne of trout at market size (product-based) or per m^2^y (surface-based).

**Table 6.** Comparison of the results assessed with the CML baseline method (Guinée, 2002) and Cumulative Energy Demand indicator (Frischknecht et al., 2007) with literature data on conventional production systems. Impacts are scaled on 1 tonne of trout.

|  | Global warming<br>(kg CO<sub>2</sub> eq.) | Acidification<br>(kg SO<sub>2</sub> eq.) | Eutrophication<br>(kg PO<sub>4</sub><sup>-</sup> eq.) | Terrestrial ecotoxicity<br>(kg 1,4 DCB eq.) | Freshwater ecotoxicity<br>(kg 1,4 DCB eq.) | Cumulative Energy<br>Demand (GJ) |
| --- | --- | --- | --- | --- | --- | --- |
| This study | 2571 | 15 | 57 | 114 | 873 | 68 |
| Literature |  |  |  |  |  |  |
| Flow-through system | 1157-3561 | 10-19 | 46-75 | 17-169 | 1290* | 30-78 |
| RAS | 2043-13622 | 13-46 | 4-21 | - | - | 63** |

Values for flow-through systems were taken from eight studies (Aubin et al., 2009; Boissy et al., 2011; Chen et al., 2015; d'Orbcastel et al., 2009; Dekamin et al., 2015; Maiolo et al., 2021; Papatryphon et al., 2004; Samuel-Fitwi et al., 2013) while values for RAS were taken from three studies (d'Orbcastel et al., 2009; Dekamin et al., 2015; Samuel-Fitwi et al., 2013).
\* Maiolo et al. (2021)
\*\*d'Orbcastel et al. (2009)

The highest environmental gains observed in organic system compared to the conventional production were for FETP (55% less in organic system): the production of one tonne of trout induced 50 kg 1,4-DCB/tonne in conventional system, but the FETP value of the organic system was noticeably lower with 33 kg 1,4-DCB/tonne. The energy requirements (CED) for producing one tonne of trout were also noticeably different between the two production systems (30% less in organic system) with values of 68 and 53 GJ/tonne in conventional and organic production, respectively. Terrestrial acidification potential (TAP) was 28% less in organic system with values of 15 and 12 kg SO_2_ eq./tonne in conventional and organic production, respectively. Differences between conventional and organic productions were less pronounced for the other impact categories. GWP showed a reduction of 12% in the organic system with, per kg of fish produced: 2602 kg CO_2_ eq./tonne were estimated in conventional system vs. 2319 kg CO_2_ eq./tonne for the organic system (Figure 7). The environmental gains through organic production were equal for LU and TETP, both of these impact categories showing a reduction of 11% in organic system while MEP is only diminished by 7% (Figure 7).

Contributions of the rearing system components (i.e., chemicals, dead biomass, energy, equipment, feeds, fry, and farm functioning) varied according to the impact category and to the production system (Figure 7). Overall, the ranking of the different contributors among the seven impact categories remained relatively constant between conventional and organic productions with the exception of chemicals, mostly driven by liquid oxygen, accounting for a non-negligible part of the environmental impacts in conventional production but not in organic production (Figure 7). Results presented in Figure 7 show that, for MEP, FEP, and WD, farm operations contributed the most to the impacts (81-84%, 90-93% and 93-98%, respectively), and the second largest contributors are either feeds for MEP and FEP (15-18% and 5-6%, respectively) or energy for WD (1-5%). For five out of nine impact categories (i.e., LU, TAP, GWP, CED and TETP), exogenous feeds were the main contributors (96-97%, 79-90%, 66-85%, 50-79% and 53%, respectively), whatever the production systems. Equipment and infrastructures are playing a significant role in the FETP and TETP impacts in the two production systems (20-53%) while their role is relatively negligible in the other impact categories. As mentioned earlier, the most remarkable difference in the contributions to environmental impacts of the two production systems concerns the role of chemicals. Indeed, chemicals include the use of antibiotics, other veterinary products and disinfectants, the use of which remains relatively constant between conventional and organic production (Table 3). On the other hand, the major difference is related to the use of liquid oxygen, included in chemicals, only in conventional production (Table 3). Thus, while chemicals represent only <2% of CED, FETP, GWP, TAP and TETP and CED in organic production, they represent between 13% and 44% of the corresponding impacts in conventional production (Figure 7).

The sensitivity analysis results indicate a linear relationship between FCR and the environmental impacts of rainbow trout farming for the nine impact categories considered in this study (Figure 8). Across most of impact categories considered, a reduction of 0.1 kg kg^-1^ in FCR leaded to a decrease of the environmental impacts decreased by 3 to 12%. Notably, the most substantial differences were observed for FEP. However, it is worth mentioning that improve feed efficiency had a negligible effect on WD, mostly linked to the water volume derived from the river and passing through the rearing structures, reducing it by less than 1%.

**Figure 8.**
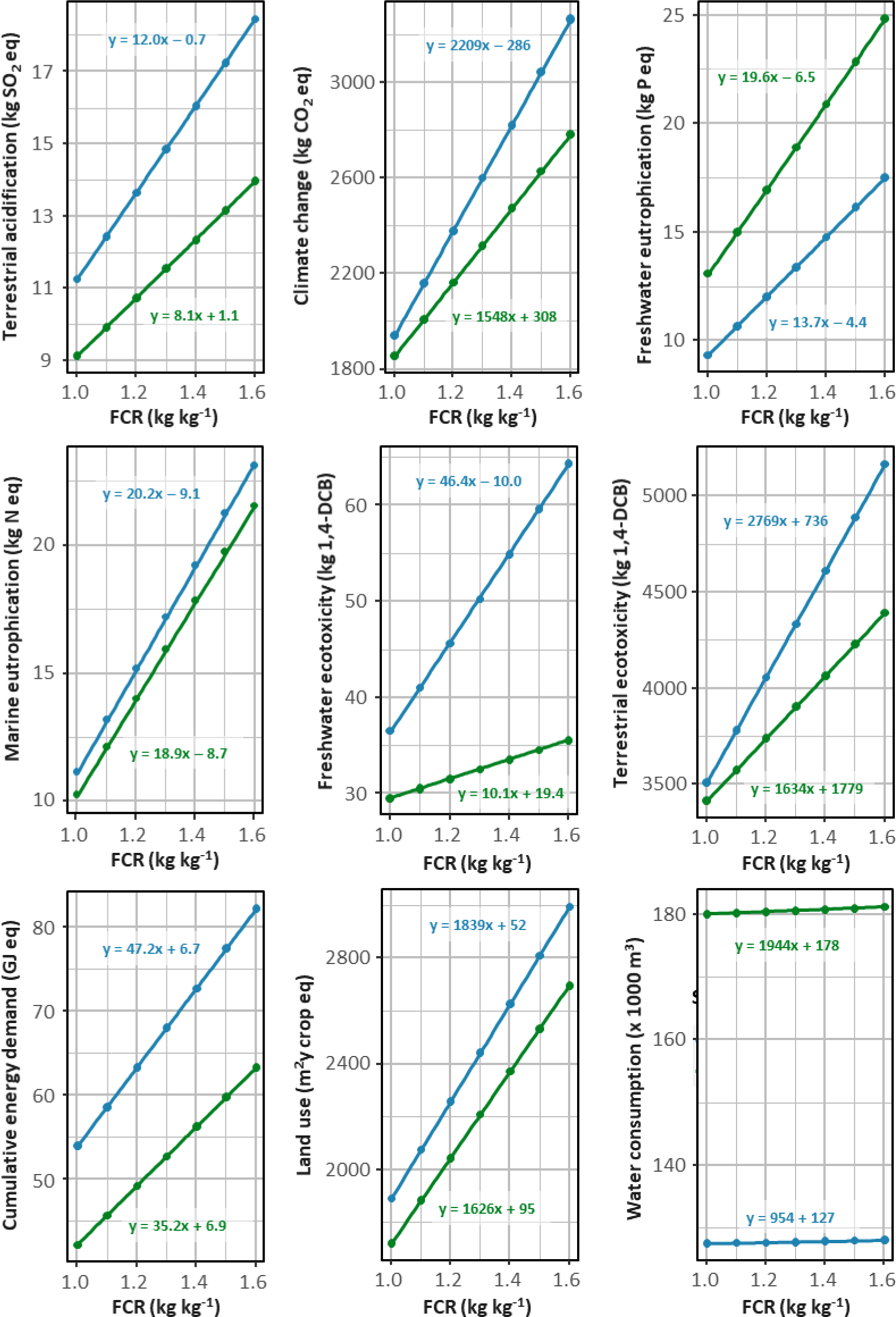
Influence of FCR variations in the environmental impacts per tonne of rainbow trout at market size in conventional and organic production system.

## 4. Discussion

Despite the rapid growth of organic agriculture production, organic finfish aquaculture remains relatively new and is still in its early stages (Mente et al., 2011). In Europe, the development of this sector has been hindered by technical challenges, such as the limited availability of organic feed and fry. Additionally, establishing effective communication strategies with clients proves difficult due to competition from other certification schemes, such as the Aquaculture Stewardship Council (ASC) or the Marine Stewardship Council (MSC) (European Comission, 2022). Furthermore, some organic farming systems experience lower yields, and previous research has suggested that the use of organic feed ingredients may lead to reduced farm eco-efficiency and increased environmental concerns (Pelletier and Tyedmers, 2007). However, there is a scarcity of peer-reviewed studies comparing the environmental impacts of conventional and organic aquaculture production systems (Biermann and Geist, 2019; Jonell and Henriksson, 2015).

The current studies in this field have predominantly followed a field-based approach, wherein data was directly collected from both conventional and organic farms to establish the LCI. However, employing such an approach may introduce certain bias, particularly regarding the distinction between differences arising from the specific production systems (conventional or organic) themselves and variations inherent to individual farming practices, which can significantly impact the interpretation of the LCA results (Chen and Corson, 2014). It is especially true in a context where the representativeness of farming practices is sometimes called into question in the LCA studies carried out in animal production (Meier et al., 2015). In this study, we employed a modelling approach, associated with LCA to compare environmental impacts of conventional and organic rainbow trout production within a hypothetical farm. The farm’s infrastructures and available surface area for production were kept constant in the two scenarios to determine the differences in environmental impacts between conventional and organic production in the same infrastructures.

Before delving into the analysis of environmental impacts between the two studied production systems (conventional and organic), it is crucial to establish a reference point by comparing the results obtained in the conventional production scenario with those from existing literature. This step allows us to compare the results from modelling with those obtained from actual fish farm data. To achieve this, we have used not only the ReCiPe method but also CML baseline method (Guinée, 2002). The latter was commonly used in previous LCA studies focusing on rainbow trout aquaculture while ReCiPe was only recently used in a rainbow trout aquaculture context notably in Italy (Maiolo et al., 2021) and Spain (Sanchez-Matos et al., 2023). Overall, the literature comparison corroborated our findings when expressed environment impact per tonne of trout (Table 6). Indeed, our results are consistent with the literature (Aubin et al., 2009; Boissy et al., 2011; Chen et al., 2015; d’Orbcastel et al., 2009; Dekamin et al., 2015; Maiolo et al., 2021; Papatryphon et al., 2004; Samuel-Fitwi et al., 2013) even if the ranges of reported values can be wide. Despite uncertainties related to varying inventory databases and CML assessment method versions, another underlying cause of the differences in environmental impacts among studies is the use of diverse production systems and varying FCRs to achieve producing the same quantity of trout (Philis et al., 2019; Sanchez-Matos et al., 2023).

The choice of functional units in LCA is a crucial point considered to conduct comparisons of production systems because its influences allocation decisions at the farm gate (Henriksson et al., 2012). Van der Werf et al. (2020) highlighted the interest in combining product-based and area-based LCA when comparing conventional and organic production systems. For instance, although organic animal production generally emits fewer pollutants per unit of land occupied than conventional agriculture (an surface-based approach), it may have higher impacts per unit of product (e.g., land occupation, eutrophication and acidification) (Meier et al., 2015). Thus, while we used one tonne of trout as a first functional unit we also expressed the environmental impacts using a surface-based functional unit (m^2^y), an original approach in LCA aquaculture studies (Bohnes et al., 2019; Pouil et al., 2023).

Overall, our study highlights a significant lower level of environmental impacts of organic production compared to conventional production. However, when impacts are expressed per tonne of trout, the WD and the FED are higher in the organic system than in conventional system. Nonetheless, it is important to be cautious when comparing the environmental performance of the two production systems using a product-based functional unit because the production capacity in the organic system is one-third lower. Specifically, the production of organic trout is limited by the lower rearing densities and reduced inputs, such as the absence of liquid oxygen (MAAP, 2010), while conventional intensive systems are managed with high stocking rates and inputs to achieve high productivity (CIPA, 2023). As a result, in the conventional production system, the environmental impacts are somewhat diluted by the larger production volume. This limitation should be considered when comparing organic and conventional systems using LCA and highlights the need to explore alternative surface-based functional units to gain a more comprehensive understanding of the comparison (van der Werf et al., 2020). By using a surface-based functional unit (m^2^y), we find that the FEP and the WD become similar between the two production systems and even slightly lower in organic system due to the absence of liquid oxygen usage. Our study demonstrates the benefits of organic trout production in terms of overall environmental impacts. Considering the nuances related to production capacity and LCA functional units is, however, crucial to gain a well-rounded perspective on the environmental performance of both systems.

The significant importance of liquid oxygen usage in conventional production becomes apparent when conducting a more detailed analysis of the contributions to environmental impacts between the two production systems. This factor often serves as the key explanation for the differences observed in impacts. Previous studies have also underscored the significance of liquid oxygen in the environmental impacts associated with aquaculture production. For instance, Song et al. (2019) highlighted that liquid oxygen contributed between 5% and 22% to all LCA impact categories. Consequently, it is evident that such production inputs should not be overlooked in LCA conducted for aquaculture production systems. Likewise, the role of aquafeeds in influencing environmental impacts is fundamental, regardless of whether it is for organic or conventional production. The importance of FCR and aquafeeds, in general, has been emphasized by numerous LCA practitioners. Several studies have already concluded that feed production constitutes a major environmental impact source (e.g., Aubin, 2013; Bohnes et al., 2019; Wilfart et al., 2023). Although organic feed helps reduce environmental impacts in many categories, its higher proportion of fishmeal and fish oil, which are rich in P (Oliva-Teles et al., 2015), leads to a greater release of phosphate into the environment in the organic production scenario, resulting in an increased risk of FEP as shown in Figure S1. It is worth noting that while feed formulations cannot be entirely disclosed due to industrial secrecy, efforts have been made to evolve these formulations. Nonetheless, these results align with the findings of Pelletier and Tyedmers (2007) who reported considerably lower environmental impacts when feeds contained reduced proportions of fish ingredients.

Given the paramount importance of feeds in determining the environmental impacts of our production systems, we investigated the effects of a change in FCR on impact categories encompassed in LCA. Our aim was to shed light on the relationship between FCR and environmental impacts per tonne of trout in our production systems. Here, we established a positive linear correlation between FCR and the environmental impacts observed. Such findings agree with previous LCA studies reporting that all environmental impacts decrease in similar proportions together with the improvement of FCR (d’Orbcastel et al., 2009; Jouannais et al., 2023; Elias Papatryphon et al., 2004). Our findings align with the conclusions drawn in a meta-analysis conducted by Philis et al. (2019), revealing a similar positive relationship between FCR and environmental impacts when comparing the environmental impacts associated with different salmonid production systems. This observation holds for changes of the FCR within a same production system and does not hold anymore across systems (Jouannais et al., 2023). Indeed, while the trend is quite clear in Recirculating Aquaculture Systems (RAS), it is notably less when considering open production systems like land-based flow-through system or open sea cages (Philis et al., 2019). Such dissimilarity can be attributed to inherent variations in the studies themselves, which become more pronounced when analysing flow-through production systems. The RAS, being more controlled, lend themselves to easier comparability across studies. In contrast, the complexities and diverse factors associated with production in flow-through systems make it challenging to draw generalizable conclusions. Nevertheless, the emergence of RAS as an alternative conventional rearing system to flow-through in trout farming has brought about new challenges, including increased energy consumption, dependence on equipment like pumps and filters, and potential greenhouse gas emissions and environmental footprint associated with energy production and waste management (Ahmed and Turchini, 2021; d’Orbcastel et al., 2009). Given the of absence of recent comparative LCA available in the literature (d’Orbcastel et al., 2009; Dekamin et al., 2015; Samuel-Fitwi et al., 2013), it could be interesting to adapt our model for comparison between flow-through systems and RAS in trout farming.

## 5. Conclusion

In our study, our modelling approach based on a hypothetical farm performing rainbow trout production under conventional and organic production constraints combined with LCA succeeded in drawing a fair description of conventional and organic scenarios of trout production enable to compare the environmental impacts at the level of the same farm. Our study demonstrates the benefits of organic trout production in terms of overall environmental impacts, which is not common regarding livestock systems. Nonetheless, our findings underscore the need for caution when interpreting LCA comparisons of such production systems, as they can be significantly impacted by methodological choices such as the chosen functional unit. Our analysis reveals that aquafeeds and liquid oxygen usage are key factors contributing to the environmental impacts of conventional and/or organic trout production systems. By recognizing and addressing the significance of these inputs, we can take further steps towards sustainable finfish aquaculture practices.

