## Supplemental Figure S1 for "Assessing the environmental impacts of conventional and organic scenarios of rainbow trout farming in France"

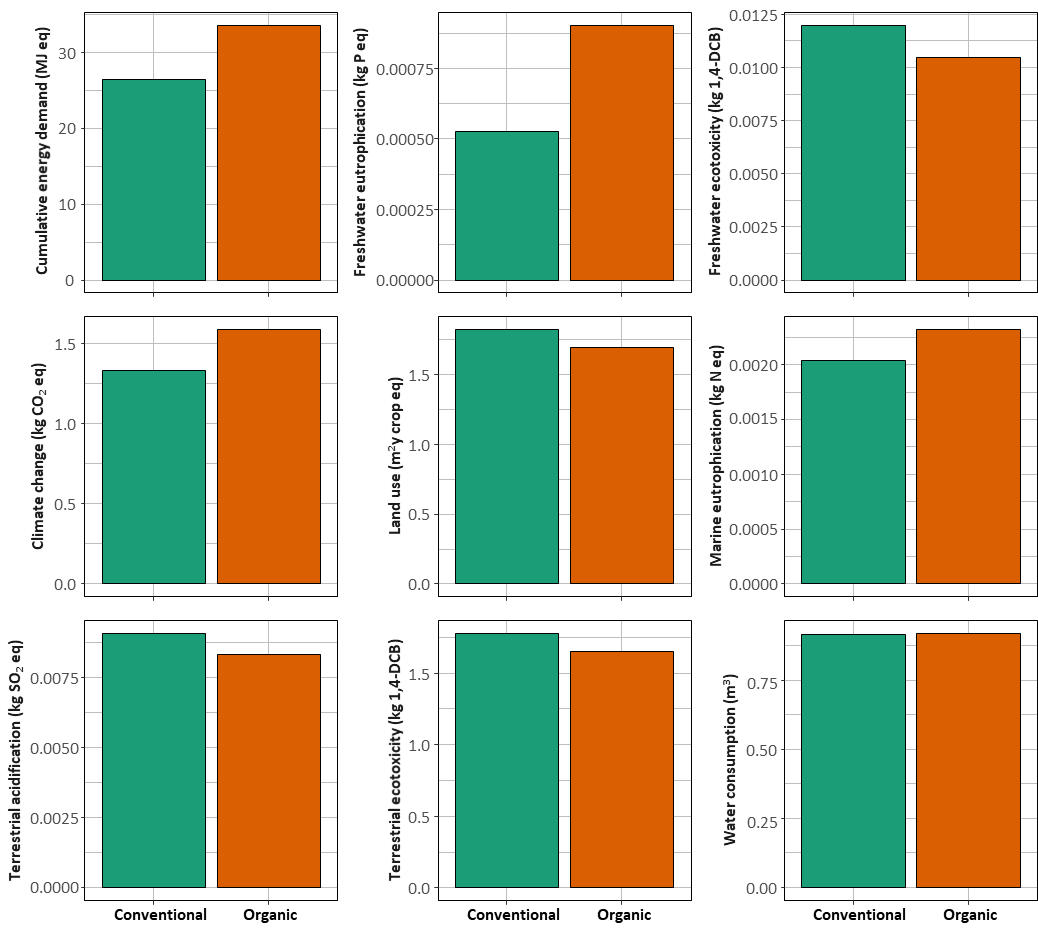

### Supplementary Material

Figure S1. Environmental impacts of conventional and organic aquafeeds used in our model for nine LCA impact category. The assessment of the impacts was carried out using ReCiPe 2016 Midpoint (H) version 1.07. Results are expressed per kg of feeds.
